# Black rhinoceros demography should be stage, not age, based

**DOI:** 10.1101/001610

**Authors:** Peter R. Law, Wayne L. Linklater

## Abstract

Biologically meaningful and standardized definitions of life stages are essential for demographic studies, especially for endangered and intensively managed species such as rhinoceros. Focusing on the black rhinoceros *Diceros bicornis*, we argue that standardized biological definitions of calf, subadult, and adult, rather than age classes, provide the appropriate basis for black rhinoceros demography. Age classes do not correlate well with the risks of mortality nor characterize, at least for females, the attainment of reproductive maturity. Black rhinoceros demography based on age classes, typical of existing literature, obscures the significance of studies of mortality and fecundity. Comparison of studies also requires standardized definitions. We propose biologically meaningful definitions of life stages appropriate for the demography of the black rhinoceros and encourage the community of rhinoceros researchers to reach consensus on standardized, demographically appropriate definitions for adoption for each rhinoceros species.

## Introduction

Age-based models of population dynamics and evolution represent a major advance over scalar models (e.g., Keyfitz and Caswell 2005, Charlesworth 1994). The difficulty of accurately aging individuals, however, motivated the introduction of age classes to which individuals could be readily assigned and the construction of stage-based models of demography (Lefkovitch 1965). These age-class stages are often defined by practical considerations (Emslie et al. 1995). But stages can be much more than just convenience in the absence of accurate age data or crude correlates of age. Stage-based models are better viewed as state-based models in which stages are well defined biological states, not necessarily or solely determined by age, with transitions between these states that are also biologically meaningful, interesting, and useful (Caswell 2001, Keyfitz and Caswell 2005, Barfield et al. 2011). Indeed, Sauer and Slade (1987a) pointed out that age-based demography is a peculiarity of human demography, not a feature of vertebrate demography in general. As noted by Caswell (2001:39), ‘For many organisms, however, the age of an individual tells little or nothing about its demographic properties.’

Size is a well known alternative state variable to age, especially for organisms with indeterminate growth (Caswell 2001:39–40 lists numerous examples including plants, fish, amphibians, and crustaceans; see also Sauer and Slade 1987a) as are discrete developmental stages in arthropods (Caswell 2001:31). Though size is sometimes employed as a surrogate for age in stage-based models (e.g., Crouse et al. 1987 for the loggerhead sea turtle *Caretta caretta*), size and age may correlate poorly. In such cases, size is not just a substitute for age but an independent state variable (Hughes 1984, Sauer and Slade 1987b); indeed, age and size, or developmental state, may each be relevant states for demography (Law 1983, Hughes and Connell 1987).

Reproductive status is another important state variable that is not always well characterized by age. Sub-adults may be characterized as individuals that are post some juvenile state (e.g., post yearling) but not yet reproductively active so that age may determine entry into the state but not exit from that state (e.g., killer whales *Orcinus orca*, Brault and Caswell 1993). In some species there may be reproductively active and non-active states that adults transition between, e.g., for birds, breeders and floaters (Hunt 1998, Penteriani et al. 2011). In various ungulate species, adult males are reproductively active when holding either harems or territories but otherwise not reproductively active and belong to male bachelor herds (Jarman 1974, Estes1991, Skinner and Chimimba 2005). In addition to the obvious difference in fecundity, survival in such cases may be state dependent.

Hence, even when ages are known, the most biologically relevant state variables to demography may not be purely age-based states. Our aim in this article is to point out that the familiar stages of calf, sub-adult, and adult are more appropriate than age for the demographic study of rhinoceros and that these stages are best defined not as age classes but as biological states. Our contention is that stages better reflect the risks of mortality than age does. In fact age may correlate poorly with these stages, allowing age-based mortality to conflate distinct risks of mortality. While females exhibit a relatively simple reproductive pattern of consecutive single calves, age does not determine first calving. Unfortunately, too little is known about male reproduction and we shall advocate, for practical reasons, an age-class definition of ‘adult male’. Genetic studies could improve knowledge of male reproductive behaviour (Garnier et al. 2001) and may in the future modify the definition of the life stage of ‘adult male’. At the present, single life stages of adult female and adult male seem demographically meaningful and biologically practical. We are aware of no evidence that a post-fecund female adult stage is necessary, e.g., as in female killer whales (Brault and Caswell 1993). A post-reproductive stage may be appropriate for males when enough is known about male reproduction.

As we shall make clear when we offer definitions in the next section, we regard the calf and subadult stages to be characterized by distinct social conditions and the adult stage to be characterized by reproductive behaviour. Thus, none of these stages are purely age classes. Although we have emphasized that stages should be biologically meaningful, they must also be identifiable in the field if they are to be useful. Without discouraging studies that will deepen knowledge of reproductive behaviour, especially for males, we will propose that a single adult stage is a practical and biologically meaningful concept for both female and male rhinoceros and will not advocate a post-reproductive stage for either sex here.

Though familiar, the notions of calf, sub-adult, and adult lack standardized definitions in the scientific literature for rhinoceros and often are defined simply as age classes. For example, the age classes of Emslie et al. (1995), refining earlier criteria of Hitchins (1970), consist of six age classes for black rhinoceros: A (≤ 5 months); B (5 months – 1 year); C (1–2 years); D (2–3.5 years); E subadult (Female 3.5–7 years; Male 3.5–8 years); F adult (Female ≥7 years; Male ≥ 8 years). In the absence of known ages, each age class is characterized by a combination of horn growth and relative body size of calf to mother so that these age classes are practical classes that can be assigned on the basis of field observation. The first four would refine the ‘calf’ stage and specify a transition from ‘calf’ to ‘subadult’ at age 3.5 years. The choice of 3.5 years appears to be based on a body size criterion. But these age classes are not suitable as life stages for demographic studies. Apart from the question of whether age classes A - D provide meaningfully distinct demographic states, a mortality in class D (2–3.5 years) is potentially ambiguous as to whether the animal was still dependent upon its mother or not. Yet this demarcation between dependence and independence upon the mother is biologically relevant for understanding the risks of mortality (Owen-Smith 1988:133). Similarly, mortality of an animal in the E class may be ambiguous as to whether it is a mortality of a calf, subadult, or adult when these terms are defined as demographic states.

We advocate the adoption of species-specific standardized definitions for each rhinoceros species of these stages as biological states so that demographic studies of each of these important species will be meaningful and comparable. Understanding the dynamics of these critically endangered species may be compromised by the use of age rather than stage and by the lack of standardized definitions. For specificity, we will focus on the black rhinoceros *Diceros bicornis.* After reviewing the scientific usage of the terms calf, sub-adult, and adult, we offer candidate definitions for these terms as life stages of the black rhinoceros. In so doing, we will make clear that these stages are not purely age based.

## Standardizing ‘Calf’, ‘Subadult’ and ‘Adult’ as Biological States

For ‘calf’ and ‘subadult’, the only issue concerns survival within these stages as, by definition, both stages will have zero fecundity. The stages reflect different risks of mortality: calves are dependent upon their mothers for food and security; subadults must provide their own food and security. Thus, the calf stage is biologically characterized as the state in which maternal investment in offspring is concentrated, whereas for the subadult stage direct maternal investment has ceased. The definitions of these stages must reflect this fundamental difference.

As noted by Owen-Smith (1988:133), ‘In most species the juvenile period ends when the young animal is driven away by the mother around the time of birth of the next progeny.’ We do not attempt to define both infant (neonate) and juvenile stages, as a distinction between total and only partial dependence upon the mother would at best be difficult to observe in the wild and so transition from infant to juvenile would have to be set by some predetermined age that would require biological motivation. Perhaps further studies of black rhinoceros might justify such a distinction. In any case, the transition from calf to subadult occurs when the calf separates from the mother; if separation is not itself observed, it may be taken to coincide with the birth of the mother’s next calf, which is more conspicuous. Even though after separation an offspring may re-associate with its mother and younger sibling (Owen-Smith 1988:136–137, Lent & Fike 2003) or even another mother-calf pair (personal observation by PRL in the Sinamatella IPZ population, Zimbabwe), we do not regard such behaviour as undermining the definition of subadult as independent of its mother or another adult. Such associations are thought to be more temporary and though they may reduce the chance of mortality, that possibility is an individual characteristic, not a stage-or age-based trait.

The definition given so far, however, is not complete as it is also known that a calf may maintain continuous association with its mother beyond actual dependency, e.g., in the absence of another birth, a failed birth, or early mortality of the younger offspring. Such behaviour is best viewed as idiosyncratic and of social significance rather than demographically informative. Mortality of the individual during such an association but at an age beyond which direct maternal investment has ended is more appropriately viewed as the death of a subadult rather than a calf. It is appropriate therefore to set an age by which the transition from calf to subadult is presumed to have taken place if separation from the mother has not occurred.

To assign such an age, we relied on Owen-Smith (1988) and Skinner and Chimimba (2005) for summaries of the older literature. For the age at which separation of calf from mother occurs, Owen-Smith (1988:136) reported an age range of 2.2–3.3 years while Skinner and Chimimba (2005:536) reported 2–4 years of age. Walpole et al. (2001) defined calves to be at most three years of age but subadults as 4–7 years of age; Hrabar and du Toit (2005) defined ‘juveniles’ to be less than three years of age and subadults to be at least three years of age; Brodie et al. (2011) distinguished between neonate calves (up to 1 year old) and ‘juveniles’ (1–4 years old). Weighing up these various assessments, we propose the calf-subadult transition should be taken to occur at separation from the mother, birth of the mother’s next calf, or on the 4^th^ birthday, whichever comes first.

We propose that ‘subadult’ means effectively independent from the mother but prior to reproductive maturity, which refers to the behavioural, not just physiological, capability of reproduction. According to Owen-Smith (1988:139), ‘Females attain adult status following the birth of their first calf….the term subadult is applied generally to the complete period from breaking of the mother-offspring bond to attainment of social maturity.’ We concur and do not attempt, for example, to define sexual maturity as the time of the first attempt at mating, but rather that of the first successful birth. Hence, we take females to transition from subadult to adult stage at their first calving. As with the calf-subadult transition, however, it makes sense to also impose an age limit on when this transition occurs, otherwise fertility rates are artificially inflated as a female that suffers delayed reproduction or fails to reproduce altogether must still be construed as adult at some point independently of its reproductive history. Owen-Smith (1988:140-141) noted considerable variation in age at first calving for black rhinoceros, with means from different areas and for captive animals occurring in the sixth year. Such variation reflects the fact that age at first calving is sensitive to resource availability in large herbivores (Gaillard et al. 2010). Skinner and Chimimba (2005:537) said subadults grow until 7–8 years of age but with no sex specific information given. Walpole et al. (2001), Hrabar and du Toit (2005) and Brodie et al. (2011) took females to mature at five years of age while Ferreira et al. (2011) adopted six years of age as the age of maturity. Emslie et al. (1995) designated females to be adults as of age 7. We suggest the transition from subadult to adult for females occurs at first calving or at the 7^th^ birthday, whichever comes first.

Hitchins and Anderson (1983) observed that spermatogenesis in male black rhinoceros in the Hluhluwe/Corridor/Umfolozi Game Reserve Complex was not observed prior to the age of eight, was evident in eight year olds, but that breeding was not observed prior to the age of nine. Skinner and Chimimba (2005) quoted these observations and further noted that successful breeding by males as young as 6.5–7 was reported elsewhere (in the absence of older males). Walpole et al. (2001) and Hrabar & du Toit (2005) both took males to be adult as of age 7 but Emslie et al. (1995) as of age eight. Brodie et al. (2011) and Ferreira et al. (2011) did not distinguish the sexes as regards the age of maturity. Until paternity can be routinely assigned in wild populations, a purely age-based criterion of maturity for males appears unavoidable. It seems prudent to follow Emslie et al. (1995) and propose that the transition from subadult to adult stage for males occurs at the 8^th^ birthday. Even if a male is not reproductively successful after reaching age eight, it would seem appropriate to assign both their mortality and fertility to the adult stage class.

In a recent paper on black rhinoceros population dynamics, Brodie et al. (2011) employed six life stages: neonate calf; male or female juvenile (1–4, which appears to mean up to age five); male or female adult (5+); mother-with-neonate. These stages were employed as states in a multi-state mark-recapture analysis and then as stages in matrix models of the population dynamics. The mother-with-neonate stage was employed following Fujiwara and Caswell (2002) because it permits estimation of fertility as a transition rate from adult female to mother-with-neonate. They ended up setting the transition rate from juvenile to mother-with-neonate to zero and the survival rate of mother-with neonate to equal that of female adults without neonates. Thus, the mother-with-neonate stage is effectively a mathematical device rather than biologically motivated and their life stages correspond in effect to the usual calf-subadult-adult life stages, albeit with age divisions reflecting the lack of fixed notions in the literature.

Brodie et al.’s paper illustrates that pragmatic considerations may motivate the employment of additional stages other than those of calf, subadult, and adult in matrix models of black rhinoceros demography; matrix models are after all *models* of demography. The most obvious advantage of age-based matrix models is that transitions between consecutive ages occur automatically and depend only upon survival. The transitions from calf to subadult and from subadult to adult, as already noted, are more biologically meaningful but more difficult to model as the probability of transition is not equal across all individuals of a given stage (calf or subadult). In the study of Law et al (2013), no calf transitioned to subadult before its first birthday (not surprising as a gestation period of 15 months permits an extended mother-calf association) but several did before their second birthday; moreover, no female subadult transitioned to adulthood during its first two years of being a subadult, but such transitions did occur during the third year. For males, by virtue of our definitions, no individual is beyond its fourth birthday upon becoming a subadult and becomes an adult upon its eighth birthday so the same fact is true by definition. Hence, to more accurately model transition probabilities, at least for the study of Law et al. (2013) but most likely for black rhinoceros in general, for matrix models one could divide the calf state into two sub-states: C_1_, for the first year of life; and C_2_, for the remainder of the calf stage; and the subadult state into three sub-states: S_1_, the first year of subadulthood; S_2_, the second year of subadulthood; S_3_, the remainder of subadulthood. Note that age of individuals in each of these sub-states of subadulthood is indeterminate. Transition from C_1_ to C_2_, from S_1_to S_2_, and from S_2_ to S_3_ occur automatically and depend only upon survival, while transition from C_2_ to S_1_ and from S_3_ to the adult state more accurately model the biological transition from calf to subadult and from subadult to adult, respectively. In line with our argument that calf, subadult, and adult are the relevant biological states for black rhinoceros demography, we would maintain that the survival rate for calves is employed for both C_1_ and C_2_ and that for subadults employed for S_1_, S_2_, and S_3_, rather than employing sub-state-specific rates. Thus, a six-‘stage’ matrix model may be constructed to model (female) black rhinoceros demography without contradicting the argument of this paper.

The Rhino Management Group’s data set includes records of females in their early thirties that had recently calved, while the longest lived black rhino in captivity, according to the 2005 international rhino studbook (Ochs 2005), was 37. Two orphaned male black rhinoceroses raised by the Sheldricks lived in protected isolation from other rhinoceros to the ages of 40 (natural death) and 38 years and seven months (speared to death) (Patton 2013).

While scant, these data suggest no need for a post-reproductive life stage for females. It would seem in any case difficult to provide a practical and reliable definition of a post-reproductive stage for female rhinoceros. For example, some six years separated the penultimate and last calves of a female in the Great Fish River Reserve; the female itself died about two-and-a-half years after the birth of its final calf, at the age of about 30 (the calf having already died within nine months of its own birth). Males, on the other hand, may require a post-reproductive life stage though it too might be difficult to diagnose. Again, much remains to be learnt regarding male reproductive success and adult survival.

## Discussion

We restate our proposed definitions in Table 1. When birth dates are unknown, to determine the ages that play a role in the definitions of life stages one can resort to the various guidelines based on body size, horn size, and for mortalities tooth eruption and wear (Emslie et al. 1995, Adcock 1997, Hitchins 1970, 1978).

**Table 1.**
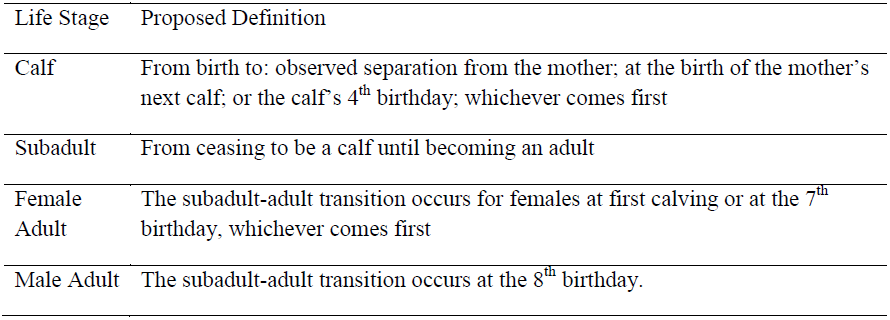
Proposed definitions of biological life stages for demography of black rhinoceros. No post-reproductive adult stages are defined but future research may require them, particularly for males. Future research may also provide a non-age-based criterion for the subadult – adult transition for males.

These definitions were employed in Law et al. (2013), a study of mortality and female reproduction for the black rhinoceros population on the Sam-Knott-Kudu-Reserve sector of the Great Fish River Reserve from the reintroduction of black rhinoceros in 1986 through December 2008. Birth dates were known for all animals born on the reserve. With these definitions, for rhinoceros born in this population the mean age (in months) that female calves became subadults was 28.0 (± 5.9; *n* = 45), that male calves became subadults was 29.5 (± 6.9; *n* = 33), and that female subadults became adults was 76.9 (± 8.3; *n* = 19). Only three calves transitioned to subadulthood by virtue of reaching the age of four years prior to separation from their mother, one female and two males. Nine female subadults transitioned to adulthood upon reaching the age of seven years without having calved; three of these females had still not calved at the end of 2008, when they were 116, 88, and 87 months of age, respectively. In this study, there was only one calf mortality and mortality was concentrated in the subadult stage (it is a young population so few animals died from old age during the study period). All the subadults that died did so after separation from their mother, underscoring the biological import of this event in rhinoceros demography rather than age.

Hrabar and du Toit (2005) reported nine non-adult deaths in their study, all of which were ‘juveniles’ (less than three years of age) but did not report whether these individuals died before or after separation from their mother. Our point of view is that this data therefore lacks important biological content and cannot, for example, be compared directly with the mortality results of Law et al. (2013) quoted in the previous paragraph. In their study, Brodie et al. (2011) found mortality to be highest for neonate calves but also nontrivial for ‘juveniles’ (1–4 year olds, which appears to mean up to five years of age). Their ‘juvenile’ mortality conflates what we have distinguished as calf and subadult mortality. Though the mortality pattern for their study is unambiguously different from the study of Law et al. (2013) the biological importance of this difference is obscured to some extent by the different notions of life stages and their biological content. In their study, Ferreira et al. (2011) found that mortality was highest for rhinoceros aged 5–6 years but that males aged 2–4 years also suffered notable mortality. Again, the biological significance of these mortalities is obscured because the age classes employed do not recognize the biologically important life event of separation of calf from mother that constitutes a state transition.

Similarly, the ‘adult’ mortalities reported by Hrabar and du Toit (2005), Brodie et al. (2011) and Ferreira et al. (2011) are ambiguous due to the purely age-based notion of female adulthood employed by these authors and likely conflates subadult and adult mortality for females. Moreover, a purely age-based designation of female adulthood may assign some fecundity to subadults and underestimate female fecundity by including individuals that are still subadults in our sense.

Defining life stages as biological states, albeit ones practical for field studies, is an essential first step in demographic studies to obtain biologically meaningful results. Standardized definitions are in any case necessary for valid comparisons amongst studies. Demographic studies of wild populations are vital for understanding and managing black rhinoceros. We believe we have offered here robust definitions of life stages for the demography of black rhinoceros. We encourage the community of rhino researchers to openly debate this issue and reach a consensus on definitions to be adopted and advocated by the African Rhino Specialist Group. A similar (though not necessarily identical) set of standardized definitions should also be adopted for white rhinoceros *Ceratotherium simum*, and at least for the Indian rhinoceros *Rhinoceros unicornis* by the Asian Rhino Specialist Group.

